# Spatially Explicit Models to Investigate Geographic Patterns in the Distribution of Forensic STRs

**DOI:** 10.1101/051375

**Authors:** Francesco Messina, Andrea Finocchio, Nejat Akar, Aphrodite Loutradis, Emmanuel I. Michalodimitrakis, Radim Brdicka, Carla Jodice, Andrea Novelletto

## Abstract

Human forensic STRs are used for individual identification but have been reported to have little power for inter-population analyses. Several methods have been developed which incorporate information on the spatial distribution of individuals to arrive at a description of the arrangement of diversity. We genotyped at 16 forensic STRs a large population sample obtained from many locations in Italy, Greece and Turkey, i.e. three countries seldom represented together in previous studies. Using spatial PCA on the full dataset, we detected patterns of population affinities in the area similar to those of genome-wide SNP and STR studies. Additionally, we devised objective criteria to reduce the overall complexity into reduced datasets. Independent spatially explicit methods applied to these latter datasets converged in showing that the extraction of information on long-to medium-range geographical trends and structuring from the overall diversity is possible. All analyses returned the picture of a background clinal variation, with regional discontinuities captured by each of the reduced datasets. These coincided with the main bodies of water, i.e. the Adriatic/Ionian and the Aegean Seas. High levels of gene flow were inferred within the main continental areas by coalescent simulations. These results are promising in a microevolutionary perspective, in view of the fast pace at which forensic data are being accumulated for many locales. It is foreseeable that this will allow the exploitation of an invaluable genotypic resource, assembled for other (forensic) purposes, to clarify important aspects in the formation of local gene pools.

## INTRODUCTION

The power of forensic STR loci for individual identification has led to the accumulation of huge amounts of data, whose interest goes beyond forensic issues and potentially relates to the gene geography of human populations and microevolutionary processes. Surveys of population samples with large arrays of STR loci (Rosenberg et al. 2002,Pemberton et al. 2013) have shown the ability of these markers to reveal many aspects of genetic structure in diverse human populations at the continental and sub-continental scales. Based on these findings different authors interpreted the arrangement of the total diversity as mainly clinal or mainly clustered, at least at the spatial scale of the examined population samples (Serre & Pääbo 2004, Rosenberg et al. 2005). The same ability for the subset of autosomal STR loci commonly used for forensic purposes is not well established. By examining two large forensic datasets Silva et al. (2012) detected a progressive reduction of diversity with increasing distance from Eastern Africa, a signature of the serial founder effect which accompanied the spread of modern humans out of Africa and beyond. However, these authors found low fixation indices, and suggested that, as far as these markers have been specifically selected to maximize the within-population (or between-subjects) variance, they carry a tiny proportion of information useful for between-population inferences. Additionally, it was suggested that the ethnic composition of population panels used in surveys with forensic and non-forensic loci may have resulted in lower fixation indexes in the former ones (Steele et al. 2014).

In recent years, several methods have been developed which, in addition to genotypic data, incorporate information on the spatial distribution of individuals within a species or a population to arrive at a description of the arrangement of diversity across that space (Guillot et al. 2005, Manel et al. 2007, Jombart 2008, Yang et al. 2012, Petkova et al. 2015, Bradburd et al. 2016). These methods represent a step forward as compared to methods that use spatial information *a posteriori*, e.g. for display purposes. In fact, to investigate spatial structures other than the most evident, a method should be spatially explicit, i.e. it should directly take spatial information into account as a component of the adjusted model or of the optimized criterion, thereby focusing on the part of the variability which is spatially structured (Jombart et al. 2008). This category of methods has been increasingly used to answer questions on gene flow, barriers, admixture and the detection of sporadic immigrants in human and non-human populations.

We report here on the typing of a large population sample obtained with a fine-grained sampling scheme involving locations in four main areas along a transect over Southern Europe and the Near East, i.e. Southern Italy, Continental Greece, the Aegean Islands and Turkey (Supplemental Fig. 1 and Supplemental Table 1). Populations from the same areas are seldom represented together in previous studies. Yet, this geographical transect, which embraces most of the North Eastern Mediterranean, is crucial for the understanding of population movements that took place over many millennia, contributing to the making of the Southern European gene pool. These include, at least: i) the entry of anatomically modern humans in Europe (Mellars 2011); ii) the spread of the Neolithic culture, which brought peoples, crops and livestock to the West, most likely as a punctuated series of mainly maritime migration episodes (Lacan et al. 2011, Rowley-Conwy 2011, Pinhasi et al. 2012, Brandt et al. 2015); iii) the expansion of the Hellenic world around the 8th century B.C.E. (e.g. King et al. 2011); iv) historical conquests in Anatolia and the Southern Balkans (Ralph & Coop 2013, Hellenthal et al. 2014).

We used 16 autosomal STR loci, included in a popular kit designed for individual identification. Thus, a feeble signal of structuring could be expected, especially in the transect here examined, which represents only a subset of the continental diversity. In this work we wanted to challenge the possibility of separating the main signal of the underlying arrangement of inter-population diversity from the background noise generated by the extreme inter-individual diversity of these markers. In this regard, we reasoned that the complex geography of the area constrained gene flow and population movements, thus requiring methods based in a geographic framework. A positive outcome in detecting spatial patterns would be promising, in view of the fast pace at which forensic data are being accumulated for many locales. This will be the basis for a description of the geographic distribution of diversity at an unprecedentedly fine scale.

## RESULTS

The final dataset comprised 1559 subjects, all typed at 16 STR loci. Overall, 288 alleles were recorded, of which 28 not present in the allelic ladder provided with the kit (overladder) and named here on the basis of their molecular weight (Supplemental Table 2). Overladder alleles were abundant at SE33, where they are known to arise also from variation outside the repeated stretch (Davis et al. 2012); we did not investigate further the molecular structure of these variants and treated them as separate alleles. Overall, 55 alleles were observed only once in the entire dataset. The number of alleles per locus varied between 9 (loci D10S1248, D16S539 and TH01) and 65 (SE33).

The screening of the entire dataset with RELPAIR confirmed that all genotypes were different from each other, as expected from the low combined probability of identity (2.35E-21) of the AmpF*l*STR^®^ NGM SElect^™^ kit (Green et al. 2013).

We wanted to assay the potential presence of null alleles by estimating their frequency at each locus in all location samples (16 × 41 = 656). In this regard, in the absence of homozygotes for the null allele, the estimated frequency depends on reduced heterozygosity as compared to the expected one. We then performed in parallel an estimation of the expected and observed heterozygosities as summarized by the index Fis. As expected, the two measures turned out to co-vary (Supplemental Fig. 2). The estimated frequencies of null alleles reached up to 0.13, often grossly exceeding the values experimentally determined by comparing assays with different primer pairs and reported at www.cstl.nist.gov/biotech/strbase/NullAlleles.htm (always <1/100 subjects; frequency <0.005). By contrast, the distribution of Fis was quite symmetrical around 0, with only a slight excess of positive values (359/656 = 55%), rarely exceeding 0.3. To check whether the diminished heterozygosity was an experimental artefact or was a real population feature, we contrasted our Fis values averaged over loci with those obtained in comparable populations with 645 STRs (Italians and Palestinians from Table S2 in Pemberton & Rosenberg 2014). Among Italians (same population background but different subjects in the two studies), Fis values of +0.0090 (95% C.I. = −0.0016 - 0.0164) and +0.0123 were obtained in the two studies, respectively. Among Palestinians (same subjects in the two studies), the Fis values were +0.0124 (95% C.I. = −0.0414 - 0.0494) and +0.0186, respectively. We then concluded that our genotyping did not suffer a significant allele loss (by either PCR failure or profile reading errors) and interpreted the reduced heterozygosity as an effect of endogamy in our closely geographically confined location samples. Therefore, we continued our analysis by considering all loci truly codominant. In these conditions, the exact test for the Hardy-Weinberg equilibrium (HWE) did not show departures from the expectation (Supplemental Fig. 3). The mean of population-specific Fis averaged over loci was 0.0119, ranging −0.058 to +0.045, with values above the average for 5 out of 6 Turkish locations, in agreement with consanguinity reports (www.consang.net), and 6 out of 7 for Continental Greece (Supplemental Fig. 4). Allele frequencies are reported in the Supplemental Table 3.

### Spatial analysis of the full dataset

The large number of alleles at the 16 loci (288), many of which with low frequencies, predicted a low signal/noise ratio in analyses of inter-population differentiation, due to inflated intra-population variances (Silva et al. 2012, Steele et al. 2014). The overall Fst among the 41 location samples was 0.0022 (P<1E-4), showing that the sampling scheme had the power of detecting genetic differentiation, though low. With such low level of differentiation, a bidimensional plot of the pairwise Fst values (nmMDS) not only did not cluster any of the locations by geography, but positioned even the outgroups within a unique cloud (not shown). However, when subjects from locations in Italy, Greece, Crete, Aegean Islands and Turkey were grouped into 11 larger samples, a pattern coherent with geography was obtained. The nmMDS plot (stress=0.183) summarizing this 13 × 13 (including the outgroups) pairwise Fst matrix (Supplemental Fig. 5) displayed the two outgroups at opposite ends on axis 1, with Continental Greek samples oriented towards the North-Western outgroup and the Turkish samples towards the South-Eastern outgroup. Italian and Cretan samples occupied a central position. The same matrix was significantly correlated (r = 0.42; P =0.017) with the matrix of geographic distances. This result was encouraging in prompting the use of spatially explicit models to highlight geographical patterns more clearly, and at the resolution of our sampling scheme (41 locations).

We first applied the Spatial PCA method implemented in adegenet (Jombart 2008) by using the full dataset of 1559 subjects × 16 loci (288 alleles). We obtained 7 and 33 positive and negative eigenvalues, respectively. The first two positive (global) eigenvalues (6.93E-3 and 4.33E-3) were remarkably larger than the third (1.87E-3) and the following ones, suggesting a boundary between strong and weak structures of the data (Jombart et al. 2008), though a test of the global spatial structure did not reject the null hypothesis (P=0.10). At any rate, sPC1 and sPC2 determined an increase in the Moran I of 10 and 3 folds, respectively, as compared to an ordinary PCA, indicating that the method could capture grater similarities between locations connected than non-connected along the network. In sPC1 (Figure 1A) the North-Western outgroup and most of the Italian locations (16/18) were identified by negative scores, whereas Greek and Turkish locations plus the South-Eastern outgroup were identified by positive scores. Two locations in continental Italy (Cilento promontory and Locri), one in continental Greece (Agrinion), one in the Aegean Sea (Khios) and one in continental Turkey (Central Anatolia) contrasted with this pattern. In sPC2 (Figure 1B) all of the Turkish locations and the South-Eastern outgroup were identified by extreme negative scores, whereas the North-Western outgroup, all of the Continental Greek and Western Cretan locations were characterized by positive scores. Eastern Crete showed moderately negative values. Italian locations were characterized by small negative (11/18) and positive (7/18) values.

**Figure 1:**
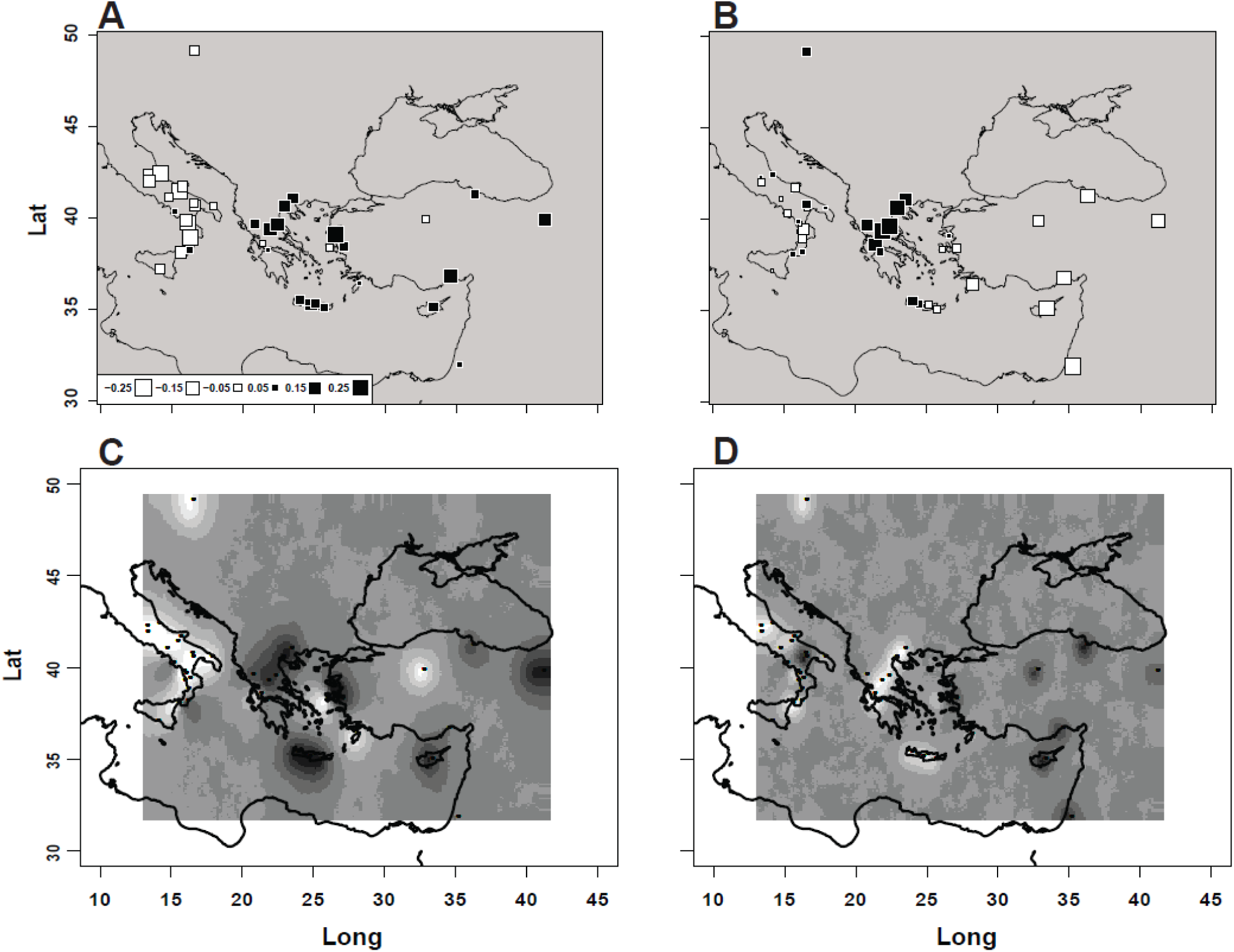
Maps of: A) scores for the 41 locations in sPC1 obtained on the full dataset with adegenet; B) scores for the 41 locations in sPC2 obtained as in A; C) posterior assignment probabilities of the 41 locations to either of two clusters obtained on the reduced dataset derived from sPC1 with Geneland; D) posterior assignment probabilities of the 41 locations to either of two clusters obtained on the reduced dataset derived from sPC2 with Geneland. In A and B white and black squares represent negative and positive scores, respectively, with square size proportional to the absolute value (inset in panel A). In each of panels C and D shades of grey indicate probabilities of assignment to one of two mutually exclusive clusters from 0 (dark grey) to 1 (white). Color versions of panels C and D are reported in Supplemental Fig. 7.

We next asked which alleles were the main contributors to these (global) patterns. We then examined the distribution of squared loadings on each of sPC’s 1 and 2, using increasing levels of stringency (Table 1). All loci were represented in the allele lists, with some of them (e.g FGA and D12S391) appearing only in the top 10% quantile. On the other hand, some loci (e.g. D10S1248 and D3S1358) contributed strongly to both sPC’s, sometimes even with the same allele. Some loci contributed to the same sPC with more than one allele, a not unexpected finding in view of the intrinsic negative correlation between the frequencies of alternative alleles at the same locus. Finally, only in 9 out of 16 loci the most frequent allele emerged, showing that a relevant spatial signal was often due to alleles that contribute only marginally to the overall variance. These observations suggested that some loci, and some of their alleles, convey a clearer signal of spatial structuring, and one or more reduced datasets can be obtained, which could retain most of the geographic information but with an abated background noise. In order to identify the optimal sets of loci and alleles we combined three criteria: a) independence between the reduced datasets derived from sPC1 and 2, i.e. the same allele should not appear in both reduced datasets; b) minimization of internal negative correlation, i.e. multiple alleles from the same locus should be avoided in the same reduced dataset; c) improved ability to detect structuring by the reduced datasets as compared to the full dataset. From the lists reported in Table 1, we thus considered optimal the choice of the alleles producing the top 2.5% of squared loadings, and recoded all individual genotypes considering each of the 8 alleles against all the remaining alleles at the respective locus (hereafter “reduced datasets”). As a result, these reduced datasets were quite complementary. First, only two loci (D2S1338 and D3S1358) contributed to both reduced datasets (each with different alleles); second, in only 3 cases two alleles of the same locus were included; third, the reduced datasets derived from sPC’s 1 and 2 produced an increase of Fst to 0.0036 (P<0.01) and 0.0031 (P<0.01), respectively (top row in Table 2), i.e. approximately 50% higher than the full dataset.

**Table 1.**
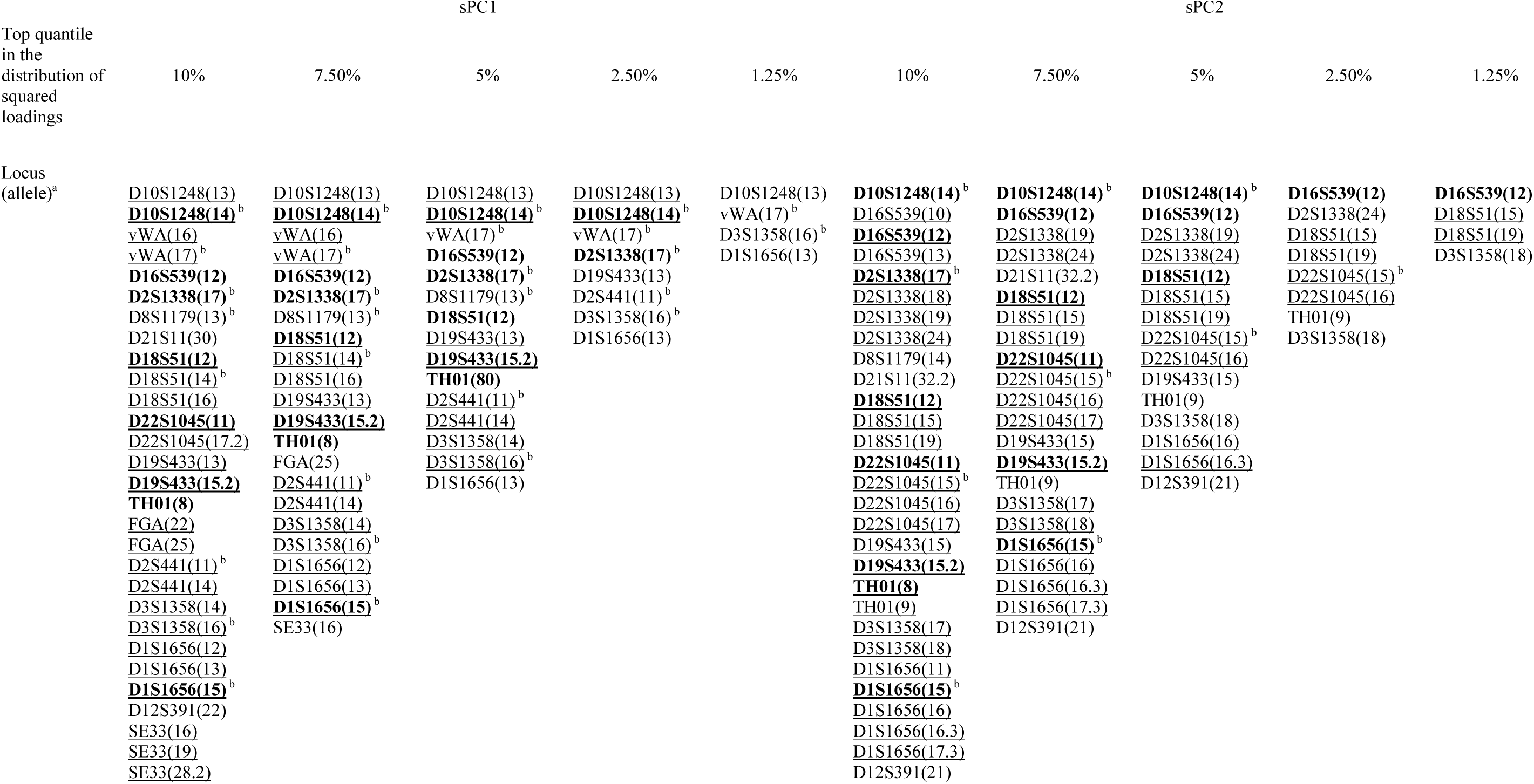
Loci and alleles with the strongest impact on the SpatialPCA eigenvalues 1 and 2. a. Listed in the order of increasing molecular weights in the blue, green, black and red series of the electropherograms b. Most frequent allele in the locus Multiple alleles from the same locus are underlined. Alleles shared between the two sPC’s are shown in boldface.

**Table 2.** Fst analysis in the main geographic regions.

| Region | sPC 1 |  | sPC 2 |  |
| --- | --- | --- | --- | --- |
|  | Fst | P | Fst | P |
| All 41 locations | 0.0036 | <0.01 | 0.0031 | <0.01 |
| Italian Peninsula | 0.0056 | <1E-4 | 0.0029 | 0.08 |
| Continental Greece | -0.0064 | n.s. | -0.0044 | n.s. |
| Crete | 0.0012 | n.s. | -0.0010 | n.s. |
| Turkey | 0.0021 | n.s. | -0.0001 | n.s. |

The frequency maps of alleles included in the reduced datasets are shown in Supplementary Figure 6. A common feature of these maps is that a general clinal variation appears, though this is evident mostly outside the polygon defined by the sampling locations, where the arrangement of equal-frequency surfaces is the result of extrapolation. The correlograms for the individual alleles displayed different patterns in the arrangement of similarity with distance. For sPC1 four correlograms were significant over the entire range: D2S1338(17) and D1S1656(13) displayed a clinal pattern over the whole range (Bonferroni corrected P<0.005 and P<0.05, respectively), whereas D10S1248(13) and D10S1248(14) displayed a minimum of the Moran I in the 800-1200 km class (Bonferroni corrected P<0.05 and P<0.001, respectively). For sPC2 only D3S1358(18) displayed a clinal pattern over the whole range (Bonferroni corrected P < 0.05), whereas D16S539(12) also showed a minimum of the Moran I in the 800-1200 km class (Bonferroni corrected P < 0.005). Allele D18S51(19) showed a strong correlation at the shortest distances but a flat pattern in the remaining distance classes. In summary, a geographic structuring of the diversity was evident, as far as 7 correlograms out of 16 rejected the null hypothesis of evenly distributed frequencies. However, we noticed that the 800-1200 km distance class included most of the comparisons between locations in Greece vs those to the West and to the East of it. In fact, when looking at the maps more carefully, it is noticeable that almost all of them display strong foci of high/low frequencies in the areas more densely covered by our locations, and such a pattern is particularly recurrent for Continental Greek locations, especially in sPC2.

### Spatial analyses of the reduced datasets

We then wanted to control that using only 8 out of 288 alleles from each sPC did not alter substantially the spatial patterns of similarity among locations. To this aim we added to our analyses two independent methods, separately for the reduced datasets derived from sPC’s 1 and 2. The membership probability maps produced by Geneland (Figure 1C,D; Supplemental Fig. 7) replicated those obtained with sPCA on the full set of alleles in many respects. The reduced dataset derived from sPC1 (Figure 1C) produced probabilities of assignment to each of two mutually exclusive population clusters ranging from 0.99:0.01 (e.g. in the Italian Peninsula) to 0.01:0.99 (Mitilini). The same 16 Italians locations identified by negative scores in Figure 1A were assigned to a single cluster with probabilities of 0.7 or higher (white shade in Figure 1C and Supplemental Fig. 7A).

The same Continental Greek, Cretan (plus Mitilini) and Turkish locations, which were identified by positive scores in Figure 1A, were assigned to the alternative cluster with probabilities of 0.7 or higher (dark grey and red shades in Figure 1C and Supplemental Fig. 7A, respectively). The reduced dataset derived from sPC2 (Figure 1D) produced probabilities ranging from 0.96:0.04 (e.g. in Continental Greece) to 0.03:0.97 (Black Sea coast). In this case all Continental Greek and Cretan locations, identified by positive scores in Figure 1B, were assigned to a single cluster with probabilities of 0.7 or higher (white shade in Figure 1D and Supplemental Fig. 7B), whereas Turkish locations plus the South-Eastern outgroup were assigned to the alternative cluster with probabilities of 0.7 or higher (dark grey and red shades in Figure 1D and Supplemental Fig. 7B, respectively). The affiliation of some locations shifted as compared with Figure 1B. In continental Italy 14 locations were now assigned to the same cluster as the Greek ones with probabilities > 0.6 and only two to the alternative cluster with probabilities >0.8, and all Cretan locations were coherently assigned to the same cluster as the Greek ones with probabilities >0.7.

Also, the effective migration surfaces analysis confirmed these broad patterns. The reduced dataset derived from sPC1 (Figure 2A) produced lower effective migration rates across the Southern Adriatic and Ionian Seas, with increased rates to the West of this belt, a pattern indicative of isolation by distance across the Aegean Sea, and again enhanced migration in the South-Eastern sector of our sampling range. In this map, the putative connection of two Italian locations (Cilento promontory and Locri) with Continental Greek locations, apparent in Figures 1A,C and Supplemental Fig. 7A, did not show up. In fact the authors of the method remark that, while a similarity of this kind could be captured by a narrow corridor, inserting such a corridor would make the overall model fit worse (Petkova et al. 2015). The reduced dataset derived from sPC2 (Figure 2B) produced effective migration rates strongly lowered along a belt stretching from the SouthEastern outgroup to Western Anatolia, shifted to the East as compared to the discontinuity in the scores and assignment probabilities of Figures 1B,D and Supplemental Fig. 7B. To the West of this belt an area including all the remaining demes displayed high migration rates (blue in Figure 2B).

**Figure 2:**
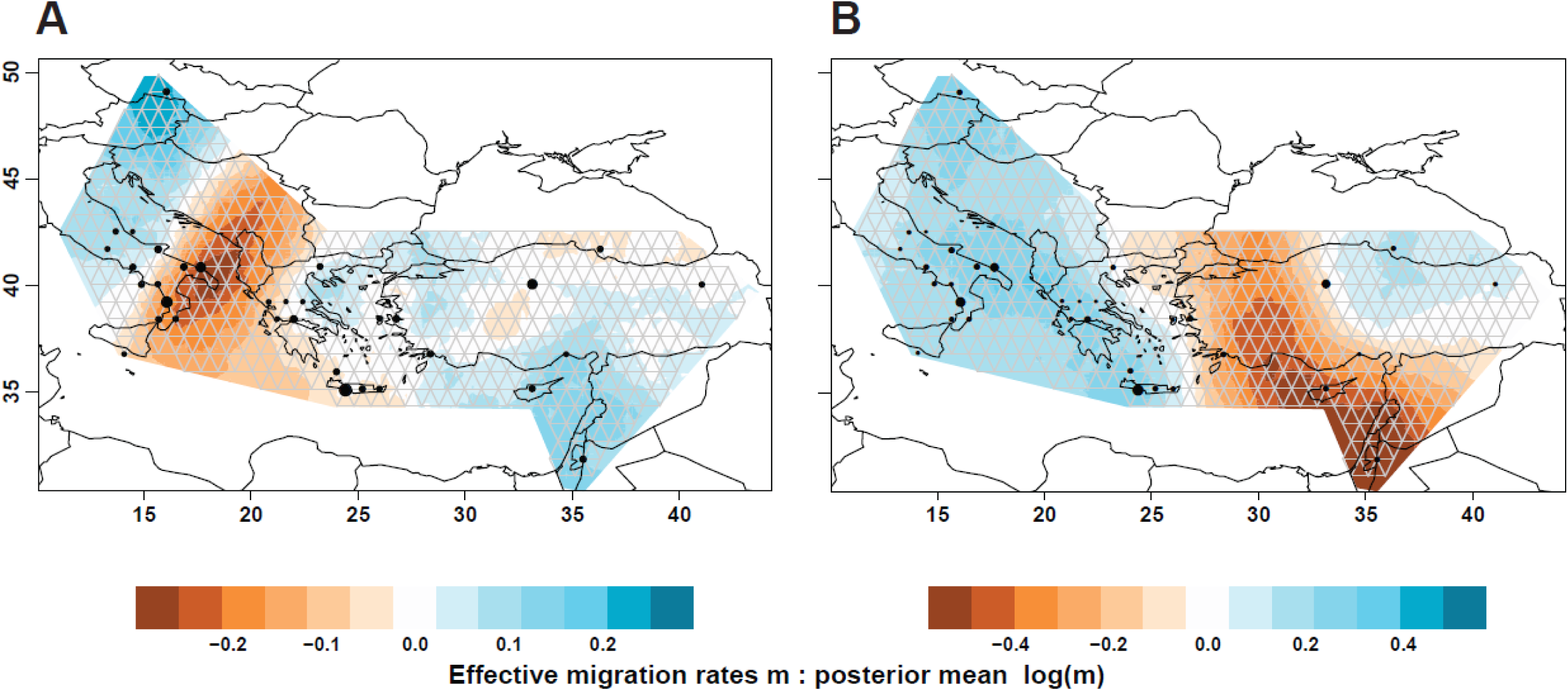
Representation of effective migration surfaces as obtained with EEMS on the reduced datasets derived from sPC 1 (A) and sPC2 (B). The coloured area covers only the user-defined polygon. The grid used by the program is shown in grey. Note that only 34 sampled demes appear (black dots, with size proportional to the n. of individuals), assigned to a grid vertex and not necessarily coinciding exactly with the original sampling location. Pooled locations were (numbered as in Suppl. Table 1): 6+7, 9+10, 13+14+15+16, 25+26, 30+32. Note the different colour scales between the two maps. In both maps brown belts correspond to low migration values, i.e. barriers to gene flow.

### Local differentiation

All the results described so far converged in showing that the extraction of information on long-to medium-range geographical trends from the overall diversity is possible. Further, a small subset of <6% of all alleles seems to retain most of the geographic information, and is able to enhance the low differentiation between locations. Analyses of both the full and the reduced datasets returned the picture of a background clinal variation onto which prominent local highs and lows generate a patchy pattern (visualized as frequency foci in contour maps). The presence of such local effects was seemingly captured also by the sPCA, which returned an excess of low magnitude negative eigenvalues (33/40), i.e. in which locations relatively close to each other (and connected in the network of Supplemental Fig. 8) produced negative Moran I’s. This raised questions on whether these minor sPC’s indeed were reporters of a real heterogeneity (Biswas et al. 2009), even between adjacent locations within the same country. We then quantified more carefully the differentiation within regional areas, by calculating the fixation index Fst (using the reduced datasets derived from sPC’s1 and 2, separately) for the locations in the areas listed in Table 2. Locations in the Italian Peninsula produced a significant Fst value with the sPC1 reduced dataset, 50% higher than that obtained on 41 locations. None of the other areas provided evidence for a similar differentiation. In pairwise comparisons, three of the Italian locations stood out i.e. Locri, Cilento promontory and Lungro. When these three locations were removed, the Fst dropped to 0.0018 (n.s.). On the basis of sPC1 scores and assignment probabilities (Figure 1A,C), Locri and Cilento promontory appeared affiliated with locations in Greece and the Aegean, which also produced positive sPC1 scores, whereas Lungro produced the most extreme negative score. In summary, with the exception of these three locations, within each of the four areas (Table 2) low differentiation was observed. This testifies of strong connectivity and prolonged gene flow between the populations that we tried to represent with our sampling, at distances ranging from some tenths (such as within Crete and Calabria) to one thousand (such as between Turkish locations) kilometers. In order to set upper and lower bounds to migration rates compatible with our observations, we used coalescent simulations, tailored on the Italian location sample sizes (Model 1 in Supplemental Fig. 9), but valid also for the other areas. When the separation of demes (14) was modelled to have occurred from 224 to 276 generations (6500-8000 years) ago, migration rates (m) lower than 0.0025 produced high Fst’s, incompatible with our observed value of 0.0018 (P<0.008 or less), whereas m values of 0.005 or greater were acceptable, with 0.01 as the best fitting (Supplemental Fig. 10A).

In this context, which factor(s) may justify the Italian peculiarity? One possible explanation is the harsh mountainous landscape of inner Southern Italy. However, we notice that the increased Fst is attributable to three locations only, distinguished from the other ones by peculiar histories. In fact, Locri and Cilento promontory coincide with two of the most important cities founded by Greek colonists in the 7-6th centuries B.C.E. (Locri Epizefiri and Velia, respectively) during the establishment of “Magna Grecia”. As opposed to other colonies of Magna Grecia, continuity of human settlement until today in the same locations or in the immediate surroundings is documented. On the other hand, Lungro was the place for the main settlement of migrants from Albania in the 15th century C.E. We then performed coalescent simulations using Model 2 (Supplemental Fig. 9), in which 3 new incoming demes join the previous 14 at 96 and 18 generations in the past. Also in this case, complete isolation could be excluded. Interestingly, however, the m value producing the best fit with the observed Fst value of 0.0056 was one order of magnitude lower than that estimated for the 14 demes (0.001 vs 0.01), and a uniform value of 0.01 for all the 17 demes turned out to be barely compatible (P=0.07) with it (Supplemental Fig. 10B). This result suggested that some degree of reduced gene flow may be responsible for the excess differentiation of the three outlier locations, in line with demographic analyses in one of them (Tagarelli et al. 2007).

## DISCUSSION

We used recently developed spatially explicit models (Guillot et al. 2005, Jombart et al. 2008, Petkova et al. 2015) to analyse forensic STR data focusing on the composition of gene pools rather than on the geographical assignment of individual genotypes. We addressed populations currently living in a geographic range of great relevance for the gene geography of Europe as a whole. Each of the four main areas here considered (Southern Italy, Continental Greece, the Aegean Islands, Turkey and sub-regions within them) can be regarded as a stepping stone for any westward migratory movement from the Levant and the associated dispersal of culture(s) (Rowley-Conwy 2011). Genetically, this is attested by phylogeographic surveys of the whole range with uniparental genetic systems (Malaspina et al. 2001, Di Giacomo et al. 2003, Cinnioglu et al. 2004, Di Giacomo et al. 2004, Semino et al. 2004, King et al. 2008, Zalloua et al. 2008, Balaresque et al. 2010, Myres et al. 2011), which led to the identification of the traces of an East-to-West gene flow, accompanied by molecular radiation of each of several lineages. Also, genetic affinities in the occurrence of male-borne lineages were observed between Aegean Islands and specific mainland populations, traceable to colonizations widely distant in time (Martinez et al. 2007, King et al. 2008). Studies directly addressed at the colonization routes led Paschou et al. (2014) and Fernández et al. (2014) to favour Crete and Cyprus (reached by seafaring) as early steps for Neolithic farmers, as compared to inland routes leading directly to the Southern Balkans. In this broad context a special case can be made of Southern Italy, which was impacted by two main immigration events from the East, i.e. the arrival of the first agriculturalists in the early Neolithic (Malone 2003) and the settlement of Greek colonists as a series of newly founded town-states starting from the 8th century B.C.E‥ The colonists in the two cases may have had different genetic ancestries and may have followed different routes. Also, they were allowed to grow for largely different amounts of time and under radically different competition/interaction regimes with already settled human groups, leading to a large uncertainty on their relative genetic legacy to the extant gene pool.

Two characteristics of the present study were challenging: first, the low informativeness of forensic STR loci for inter-population analyses even in distantly related groups (Silva et al. 2012, Steele et al. 2014) and, second, our geographic range, stretched over 30 degrees of longitude but compressed into 10 degrees of latitude. Nevertheless, some of our results qualitatively match those obtained on widely dispersed population samples, with far more STR loci or SNPs.

As to genotyping, our results revealed a background level of inbreeding in the majority of locations (roughly two thirds), which is not attributable to the use of 16 loci only, but is consistently replicated with larger numbers of loci in many world populations (see Figure S2 in Pemberton & Rosenberg 2014). Our estimated values of the inbreeding coefficient agree with the limited population size of the sampled locations and the traditional marital habits (Tagarelli et al. 2007). Also, they exceeded the values (averaged over subjects) of the direct estimations of homozygosity obtained in two population samples overlapping with those examined here, i.e. Palestinians (Pemberton & Rosenberg 2014) and Italians (Gazal et al. 2015). This indicates that, when considering local samples, a slight excess of homozygosity is a common occurrence, even in individuals more remotely related than currently detectable with closely spaced genetic markers.

The spatial analyses here performed showed that the extraction of information on geographical trends from the overall diversity is possible, provided that explicit models are employed. Additionally, we devised objective criteria to reduce the overall complexity into two reduced datasets of 8 + 8 alleles, which retained the signal of spatial trends and of genetic structuring. The most prominent feature, confirmed by three independent methods, is the detection of discontinuities in the similarity of our location samples which coincide with the main bodies of water. An Adriatic/Ionian discontinuity (Figures 1A,C and 2A) was captured by sPC1 and the reduced dataset derived from it. An Aegean discontinuity (Figures 1B,D and 2B) was captured by sPC2 and the reduced dataset derived from it. Together with coalescent simulations, this indicated that high connectivity maintained a stronger homogeneity within the geographic peninsulas and islands (through land contacts) than between them, despite documented contacts by maritime routes over millennia. As outlined above, the geographic range here surveyed was suitable to reveal trends only in the East-West axis, with little or no power to detect similarities/discontinuities on the South-North axis. Moreover, our datasets are enriched in Greek subjects as compared to other studies in the area. As discussed elsewhere (Lazaridis et al. 2014), these two factors determine that, eigenvalues 1 and 2 of sPCA cannot be expected to necessarily have a direct correspondence with eigenvalues 1 and 2 in ordinary PCA’s of studies encompassing a broader range of European populations. In these conditions, the orthogonality (Novembre & Stephens 2008) between the first two sPC’s of the present study, which we tried to maintain also in the reduced datasets derived from them, may have enhanced the Adriatic discontinuity. Also, it is possible that our locations in Continental Italy spanned a barrier across the Peninsula observed in other studies (Nelis et al. 2009, Di Gaetano et al. 2012). In this context, subtle differences emerge from genome-wide SNP studies which included Italian, Greek and a small number of Turkish subjects, concerning the affinities among the three groups. Their centroids turned out to be equidistant in the PC1-PC2 space in three studies based on the POPRES resource (Novembre et al. 2008, Wang et al. 2010, Wang et al. 2012). In none of these studies Italian and Greek subjects were so well geographically referenced as we have done here, and it is thus not surprising that in the same studies the clouds of individual points largely overlapped. However, by analysing the pattern of sharing of genomic blocks identical by descent in the same resource (Ralph & Coop 2013), an unusually little common ancestry was found between the Italian peninsula and other locations, seemingly deriving from longer ago than 2,500 years.

On the other hand, our sPC2 and the reduced dataset derived from it set the Turkish locations apart from the ones to the West, with clearcut sPCA scores, assignment probabilities and a migration barrier. This feature and the spatial orientation of this sPC establish a straightforward similarity with other analyses of genome-wide SNP data in which Italian and Turkish subjects fall into separate clinal series covering the Middle East/Caucasus and Sardinia/Continental Europe, respectively (Behar et al. 2010, Yunusbayev et al. 2011). Also, in a study based on an independent sampling as compared to those mentioned above (Lazaridis et al. 2014), the cloud of Turkish subjects was neatly separated from a poorly structured cloud of Greek/Sicilian/Continental Italian subjects. Finally, in a study using a closely spaced sampling scheme similar to the present one and network analysis (Paschou et al. 2014), subjects from Cappadocia (Central-Eastern Turkey) were separated from Greek, Cretan, Sicilian and Italian samples.

A complementary and not necessarily alternative interpretation of the global spatial patterns highlighted here is a distinctiveness of Continental Greek locations, and especially the Eastern ones, which extended to Crete. Two studies (Ralph & Coop 2013, Hellenthal et al. 2014) stressed the relevance of expansions and admixture events in the 4-10th century C.E., which shaped the ancestry of Southern Balkans up to the Aegean coasts of Greece.

Yet another aspect is the retention of information on short-scale variation in the reduced datasets. Contrasting patterns in the distribution of Y-chromosomal and mtDNA markers emerged within Southern Italy (Di Giacomo et al. 2003, Capelli et al. 2007, Sarno et al. 2014). Genome-wide SNP data did not detect an obvious geographic patterning within Southern Italy (Di Gaetano et al. 2012), in line with the lower expected fluctuations for frequencies and clines of autosomal alleles. Our coalescent simulations indicated that Fst values of the magnitude observed here could be better explained by high levels of gene flow prolonged over thousands of years. The estimated migration rates exceeded those obtained with a similar method for males and females in Asian patrilocal societies (Hamilton et al. 2005), and approach those experimentally measured in contemporary European populations (Cavalli-Sforza & Bodmer 1971, Fix 1999).

Some instances of the observed short-scale discontinuities of sPCA scores and assignment probabilities are also compatible with the historical accounts on the settlement of some of the locations in Southern Italy, which is in turn reflected also in the occurrence of Greek surnames (Fig 5.7.4 in Cavalli-Sforza et al. 1994). sPC scores and assignment probabilities for two Italian locations bear similarity to those of Eastern Greece and around the Aegean Sea, but the causal relation between the settlement of Magna Grecia and this observation remains to be determined. For example, traces of the Hellenic colonization were detected in some Sicilian locations only (Di Gaetano et al. 2008), and not in Southern Continental Italy (Tofanelli et al. 2016). Nevertheless, the method outlined here may be helpful to orient heuristic searches of specific markers in these locations, that could confirm/dismiss hypotheses on genetic contributions from the Early Neolithic Levant, the Hellenic world or the Balkans.

Pioneering studies with the Y chromosome (Roewer et al. 2005) have shown that collating forensic data can result in a high power of detecting spatial patterns even at the sub-national level. The results reported here call for a replication of the analytical procedures on additional forensic STR datasets, in order to test their performance against known features of the gene geography of Europe and other areas. These include e.g., the continent-wide SouthEast-to-NorthWest cline extending to Scandinavia and the British Isles initially described with classical pre-molecular markers (Cavalli-Sforza et al. 1994), as well as features of local populations. It is likely that enlarging or shifting the surveyed geographic range will lead to different loci and allele lists as compared to those reported here, and to different thresholds to select the most informative ones. Also, different standardized multiplex assays allow the genotyping of other sets of loci for individual identification. Intersecting the results obtained with them will come at the cost of reducing the number of shared loci but with the advantage of enormously increasing the sample size.

## CONCLUSION

Spatially explicit models empower the analysis of geographic structuring of extant human diversity at forensic STR loci up to a sub-continental and, possibly, to an even finer scale. It is foreseeable that this will allow the exploitation of an invaluable genotypic resource, assembled for other (forensic) purposes, to clarify important aspects in the formation of local gene pools.

## MATERIALS AND METHODS

### The samples

All samples were from collections of the authors, assembled in the 1980’s, 1990’s and 2000’s. The original sampling was performed by colleagues and operators at a number of collaborating Institutions and included the recording of the subject’s place of residence. The subject was also asked to report the origin of his/her parents. Recent immigrants were excluded. A total of 40 villages/towns (hereafter “locations”) were sampled (Supplemental Table 1 and Supplemental Figure 1). In most cases a sample representing a location consisted of subjects with that residence, but in a minority of cases information on residence was collected at a finer scale, and residents in the neighbourhood were assigned to the nearest location.

As far as the proposed research did not involve any issue relevant for the donor’s health, only a subset of the WMA Declaration of Helsinki and COE Oviedo Convention prescriptions were applicable and obeyed. For these reasons written consent was requested in most cases but, in some series collected before 1995, oral consent was considered sufficient and simply recorded in the corresponding log sheets (filed at the collecting Institutions). In all cases the consent included also storage and future use of the sample. Anonymized blood or DNA samples were then received at the corresponding author’s laboratory. The study was prospectively examined and approved on November 21, 2014 by the intramural ethical committee (document number 0025422/2014), who expressly considered the list of collaborators, anonymity of samples and the compliance with consent regulations of previous publications which included the same samples.

The 40 locations comprised a group of residents in Moravia (Czech Republic) (Luca et al. 2007) as a reference for the Central European population (North-Western outgroup). In order to have an appropriate counterpart, we also included in the study (41st location) the Palestinians of the CEPH HGDP panel (Cann et al. 2002), as a South-Eastern outgroup.

### STR profiles and quality controls

We obtained the genetic profiles at 16 autosomal STRs on 1984 individual assays using the AmpF*l*STR^®^ NGM SElect^™^ PCR Amplification Kit (Life Technologies inc.) under the conditions recommended by the manufacturer, in 96-wells microtiter plates. PCR products were separated in a ABI PRISM^®^ 3100-*Avant*^™^ Genetic Analyzer. All plates included a negative (water) control, 8 replicates of the reference allelic ladder provided with the kit, as well the positive control provided by the manufacturer (Control DNA *007*) and our internal control. Electropherograms were generated with the GeneMapper ^®^ ID-X software, with allele nomenclature given in number of repeats at each locus. Profiles were inspected by two independent operators. Independent spreadsheets were produced and compared. All discrepant results underwent a first round of reviewing. Profiles with missing amplification at one or more loci were discarded.

More than 300 subjects, spread over separate plates, were typed two or more times with 100% repeatability. As far as we analysed the CEPH Palestinians, we could directly compare our results with those already obtained in the same individuals (Pemberton et al. 2013). Identical genotypes were obtained for the 4 loci shared between the two studies (D10S1248, D16S539, D19S433 and D22S1045), after translating allele sizes from bp to repeat numbers.

We also used the program RELPAIR (Epstein et al. 2000) to detect hidden relatedness, with allele frequencies obtained in the whole series. Thresholds for the likelihood ratio took into account the number of pairwise comparisons within each location sample. This step led to the exclusion of 19 subjects, i.e. one member of each relative pair (Parent-Offspring = 12; Full Sibs=7). Among full sibs, the relatedness of two Palestinians (HGDP00694 and HGDP00695) was detected, as previously reported (Rosenberg 2006).

### Data analysis

#### Standard diversity indices

The potential presence of null alleles was checked with the program FreeNA (Chapuis & Estoup 2007). Allele frequencies and fixation indexes were obtained with Arlequin v. 3.5.2.2 (Excoffier & Lischer 2010), both considering (for Fis) and not considering (for Fst) the individual level. The exact test for the HWE was performed with the same program and 1 million steps in the Markov chain. The matrix of pairwise Fst values was represented by non-metric Multi Dimensional Scaling (nmMDS) with PAST v. 3.06 (Hammer et al. 2001). The Mantel test as implemented in PASSaGE 2 (Rosenberg 2001) was used to test the correlation between the Fst and geographic distance (great circle) matrices.

#### Spatially explicit models

Spatial principal component analysis was performed with the R package adegenet (Jombart 2008). This method uses the frequencies of alleles at all loci and performs a principal component analysis which takes into account simultaneously the frequency variance across locations and the similarity between locations connected in a spatial network as contrasted to those not connected (summarized by the Moran I index). It thus returns positive (and negative) scores according to whether closely spaced locations display higher (lower) similarities, thus distinguishing between global vs local frequency variation patterns. Geographic coordinates were those reported in Supplemental Table 1 and a nearest neighbour (n=12) connection network was used (Supplemental Fig. 8). The distributions of squared loadings for the 288 total alleles on sPC’s 1 and 2 were used to identify the strongest contributors to each sPC, represented in the top 1.25%, 2.50%, 5.0%, 7.5% and 10% quantiles, corresponding to 4, 8, 15, 22 and 29 alleles, respectively. Surface maps of the frequencies of the selected alleles (8 in each case, see Results) were constructed with the Kriging algorithm as implemented in the R package “fields”, using a 200 × 200 rectangular grid spanning from the minimum to the maximum longitude and latitude values. The advantage of this method is that, at each sampling location, it returns the observed value (in this case the allele frequency) and thus the density of equal-frequency surfaces reflects the steepness of the expected cline(s) along lines connecting sampled locations. Coast contours and markers for sampling locations were overlaid with R functions. Correlograms of allele frequencies were obtained with PASSaGE 2 (Rosenberg 2001), using 6 distance classes (Upper bounds: 200, 400, 800, 1200, 1600 and 3200 km), chosen to obtain an appropriate number of observations within each class, and to render the shorter distances within than between each of the three main sampling countries (Italy, Greece and Turkey). Significance was tested with 10,000 permutations.

In order to retain most of the spatially informative aspects of the data while abating the background noise as compared to sPCA, we generated genotype datasets (reduced datates) derived from sPC’s 1 and 2. In each of them we recoded the individual genotypes considering the 8 alleles with the highest squared loadings on sPC’s 1 and 2, respectively, against all the remaining alleles at the same locus (Patterson et al. 2006). Two independent spatially-explicit methods were applied to both datasets to describe affinities between location samples and patterns on anisotropic migration (barriers to/enhancements of migration).

The ability of the reduced datasets in detecting affinities between locations was assayed with the clustering algorithm implemented in the program Geneland (Guillot et al. 2005). This method employs a Bayesian approach that takes into account the spatial coordinates (in this case identical for all individuals of the same sampling location) of each genotyped individual to assign him/her to a population cluster among a number of clusters that can be bounded by the user. Both reduced datasets were then analysed under the uncorrelated model with 200,000 iterations with a thinning of 100 and an initial 20% burn-in, asking for the probability of assignment to either of two clusters (to directly compare with the qualitative result of sPCA). Such probabilities, obtained for each square of a 200 × 200 grid, were plotted from the output files with R functions as above.

In order to visualize estimated migration rates we used the program EEMS (Petkova et al. 2015). This program uses a regular triangular grid covering a polygon which embraces the entire geographic range of sampling. Each individual (in our case all individuals of the same sampling location) is assigned to the nearest vertex of the grid and the migration parameter m is estimated by Bayesian inference for every edge of the grid. The processed output consists in maps in which colours of the estimated effective migration surface correspond to local deviations from isolation by distance: in particular effective migration is low in geographic regions where genetic similarity decays quickly. We considered a polygon extending from the North-Western to the South-Eastern outgroup and spanning over most of the Italian peninsula, the western Balkan peninsula and almost the entirety of Turkey (11 to 44 degrees of longitude and 50 to 30 of latitude). We performed a number of test runs to optimize the performance of the MCMC chain and to obtain a grid which could spatially resolve most of our locations. Eventually, we used a grid of 466 triangles (demes) which resulted in the reduction of the 41 location samples to 34 sampled demes, some locations being assigned to the same vertex and pooled. In order to use relaxed priors, the variances of the proposal distributions were increased 8x as compared to the defaults. Two independent runs of 2,000,000 steps, 50% burn-in and a thinning of 5000 were used for each reduced dataset. Postprocessing was as recommended by the authors.

#### Coalescent simulations

In order to test the observed fixation indices against neutral evolutionary scenarios, coalescent simulations were obtained with Fastsimcoal2 (Excoffier & Foll 2011). We used two demographic models modified from that of Capocasa et al. (2013) and graphically outlined in Supplemental Fig. 9. In Model 1, a number of populations (initially set at 14 to simulate Italian locations) split from a large pool (100,000 current gene copies) which has been growing at a rate of 0.020/generation, i.e. a value valid also for pre-agricultural societies (Ammerman & Cavalli-Sforza 1984, Boone 2002, Hamilton et al. 2009). Generation time was assumed to be 29 years (Fenner 2005). Splitting times (in generations before present) were sampled from a UNIFORM(224:276) distribution, to account for the long time of the spread of the Neolithic cultural package in the Italian peninsula (Malone 2003), and deme sizes (in gene copies) were sampled from a UNIFORM(800:2400) distribution, to account for the autosomal effective size and larger census size as compared to Capocasa et al. (2013). These demes were set to grow at a rate of 0.017/generation, continuing to exchange gene copies with the main pool at rates of 1.0E-3 and 1.0E-4 for sending and receiving, respectively. The resulting distribution of the fixation index Fst was obtained for migration rates (m) between the 14 demes of 0, 0.0005, 0.001, 0.0025, 0.005, 0.01 and 0.02 (corresponding approximately to 0-30 migrant copies (M)/generation). A set of microsatellite loci with constrained number of alleles were modeled, to replicate the binarized recoded loci described above. For each condition 2,000 simulations, with samples of size equal to the real samples, were run and analyzed with Arlequin v. 3.5.2.2. In Model 2, three additional demes were added to the scenario with m = 0.01, joining the set of 14 at 96 (2) and 18 (1) generations ago, respectively. These latter demes were simulated to exchange gene copies with the previous 14 at rates of 0, 0.001, 0.0025, 0.005 and 0.01.

## ACKNOWLEDGMENT

We thank all anonymous donors for their voluntary participation in this study. We thank all collaborators who performed the interviews, filed them and collected the biological samples. This work was supported by grants of the Italian Ministry of Justice (grant number CUP E81J10001270005) to C.J. and of the Italian Ministry of Education (grant number PRIN-MIUR 2012JA4BTY) to A.N‥ F.M. was supported by a fellowship from the Italian Ministry of Education.

---

**Supplemental Fig. 1.**
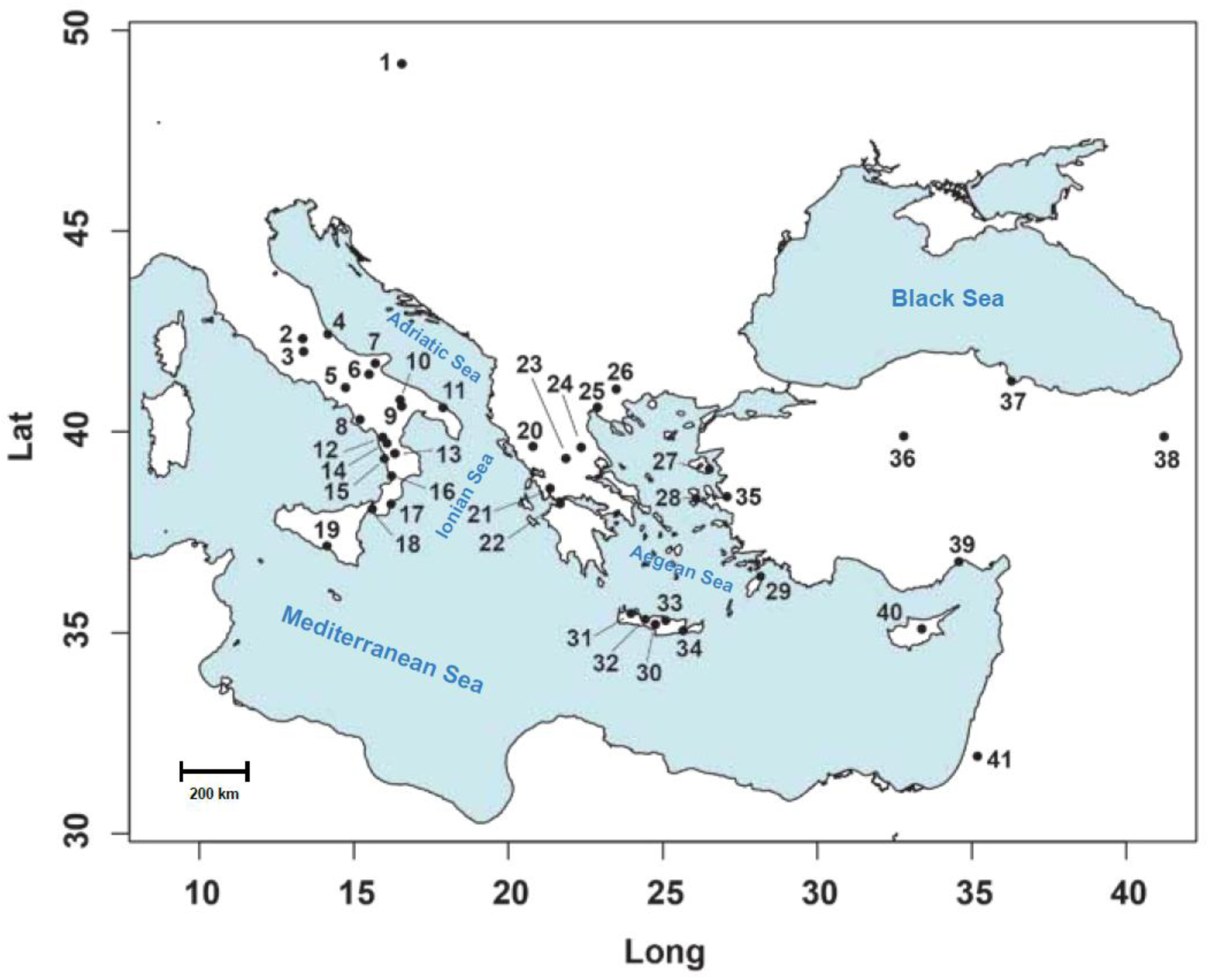
Map of Southern Europe/Northeastern Mediterranean Sea, with the positioning of sampling locations numbered as in Supplemental Table 1.

**Supplemental Fig. 2.**
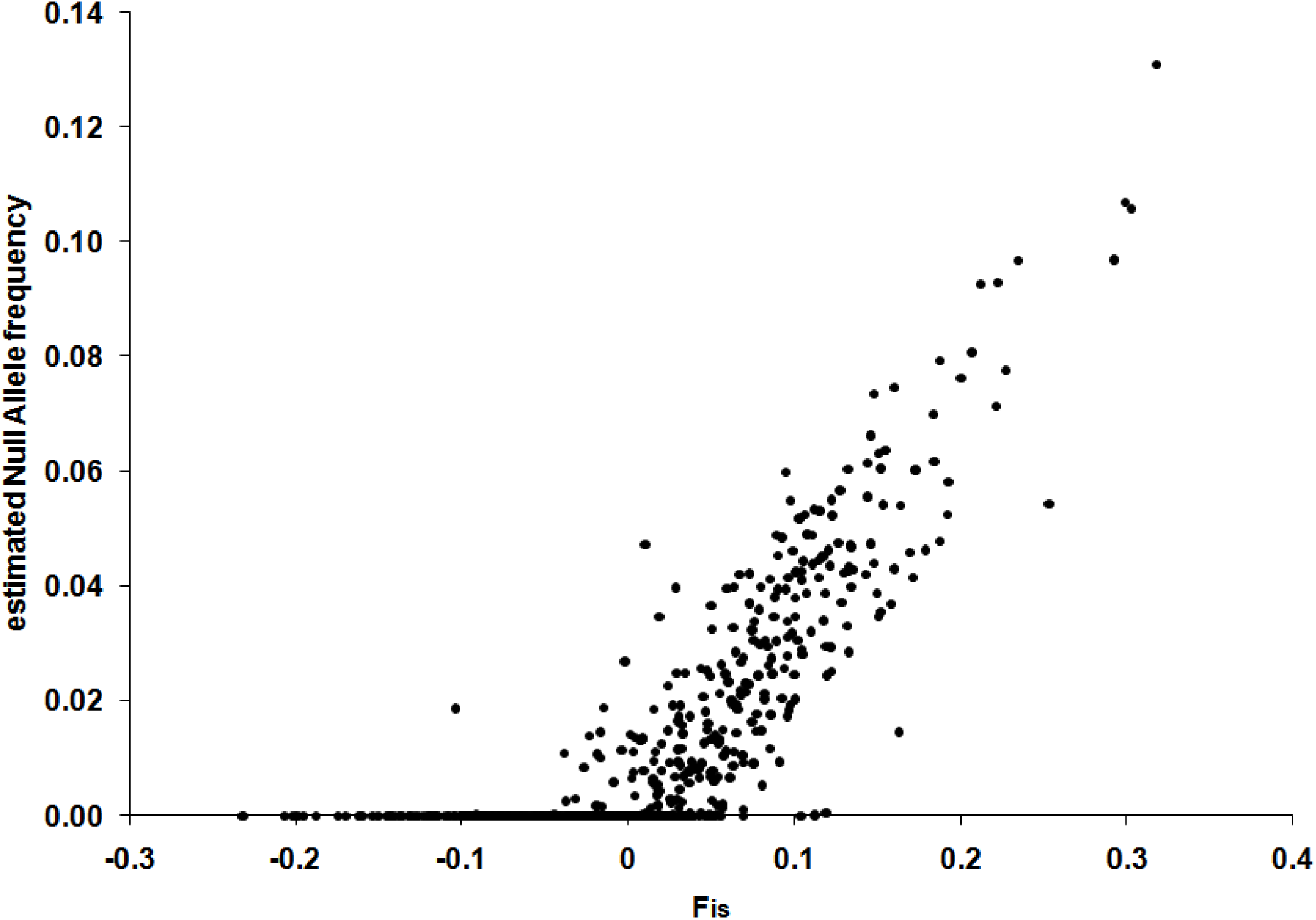
Scatterplot of Fis vs the estimated frequency of null alleles at 16 loci × 41 location samples

**Supplemental Fig. 3.**
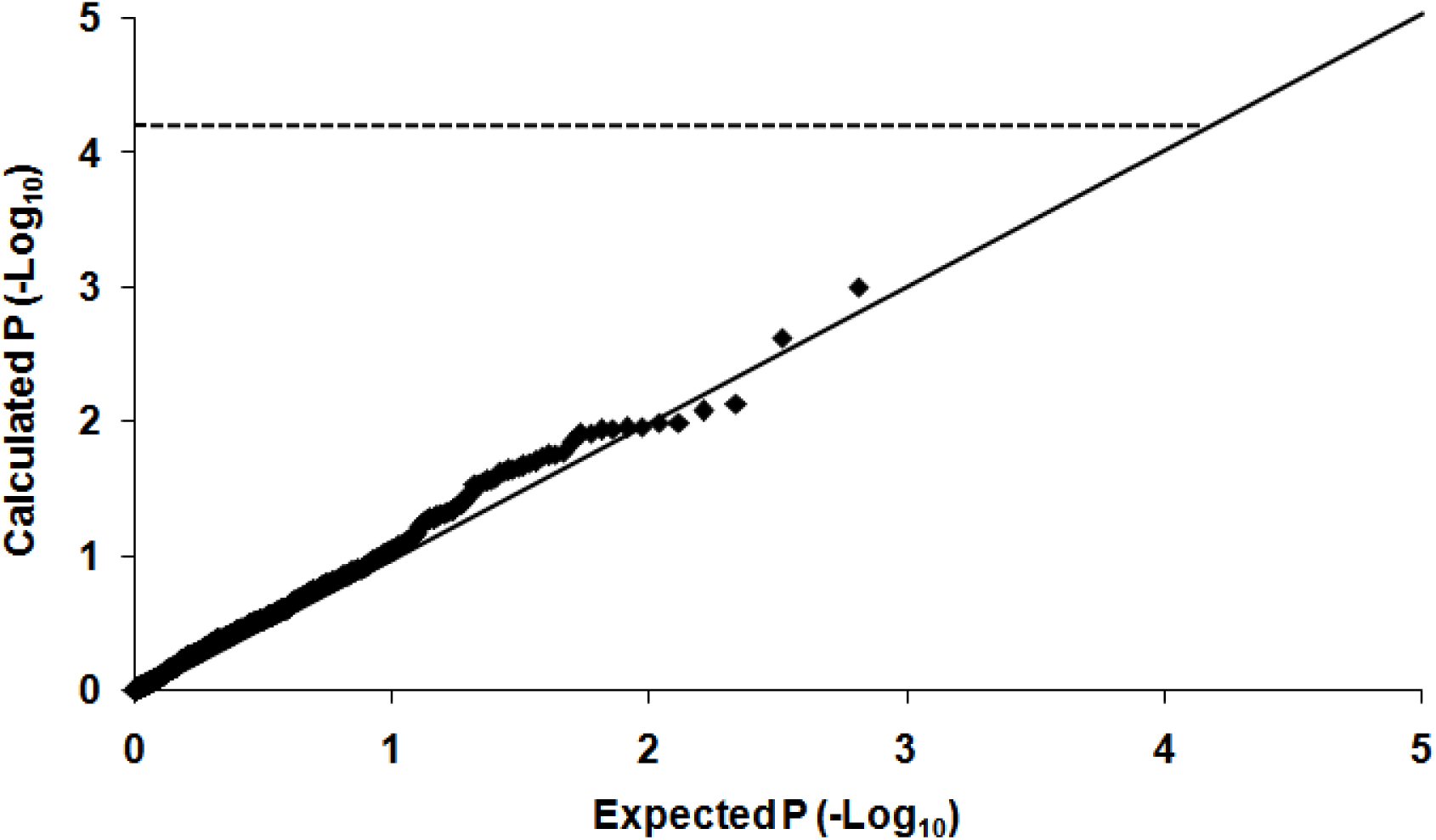
Quantile-quantile plot of the probability values in 656 tests for HWE (16 loci × 41 location samples). The solid line indicates identity between calculated and espected values. The dotted line represents the significance level (nominal α = 0.05) after Bonferroni correction.

**Supplemental Fig. 4.**
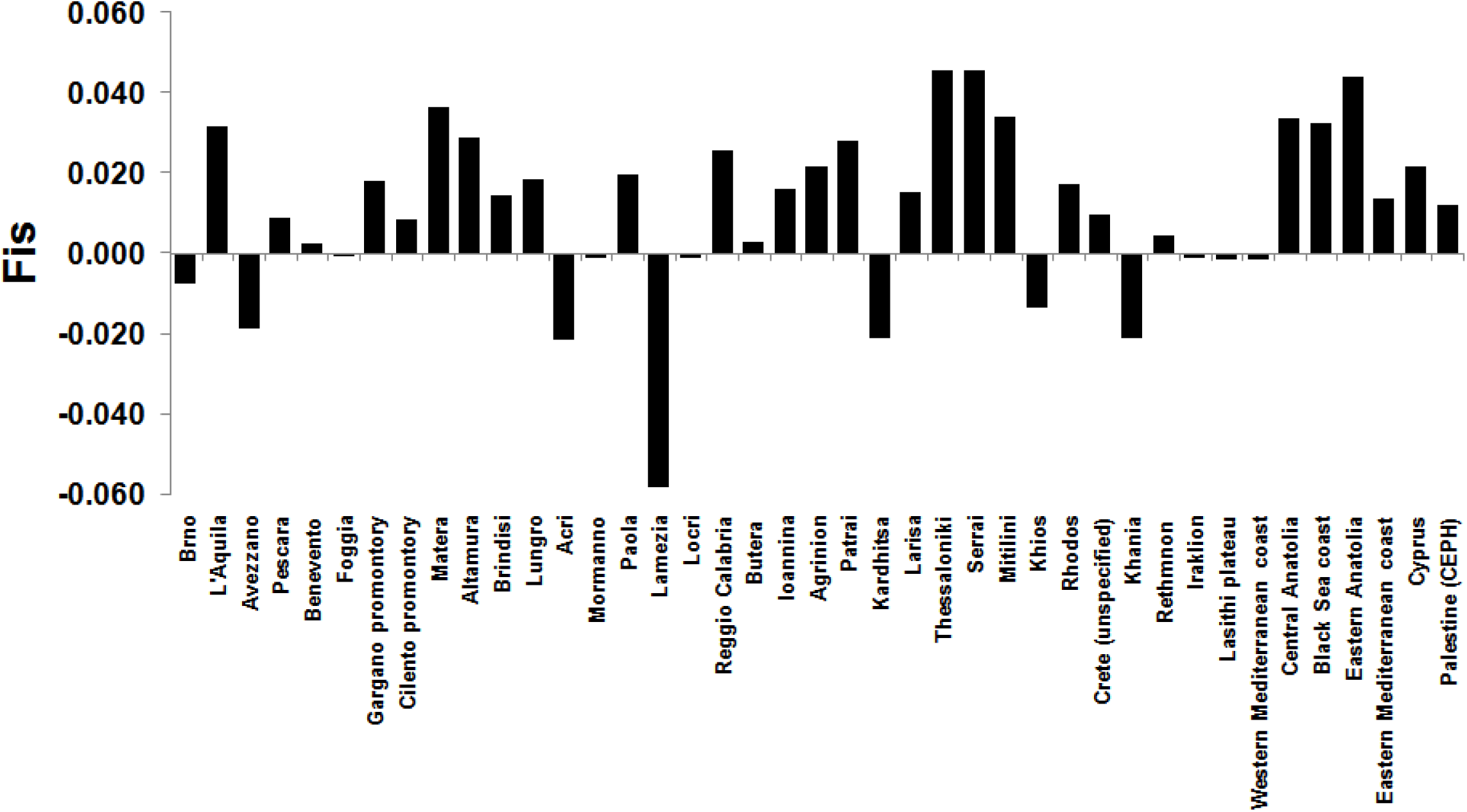
Barplot of the Fis values (averaged over loci) for the 41 locations.

**Supplemental Fig. 5.**
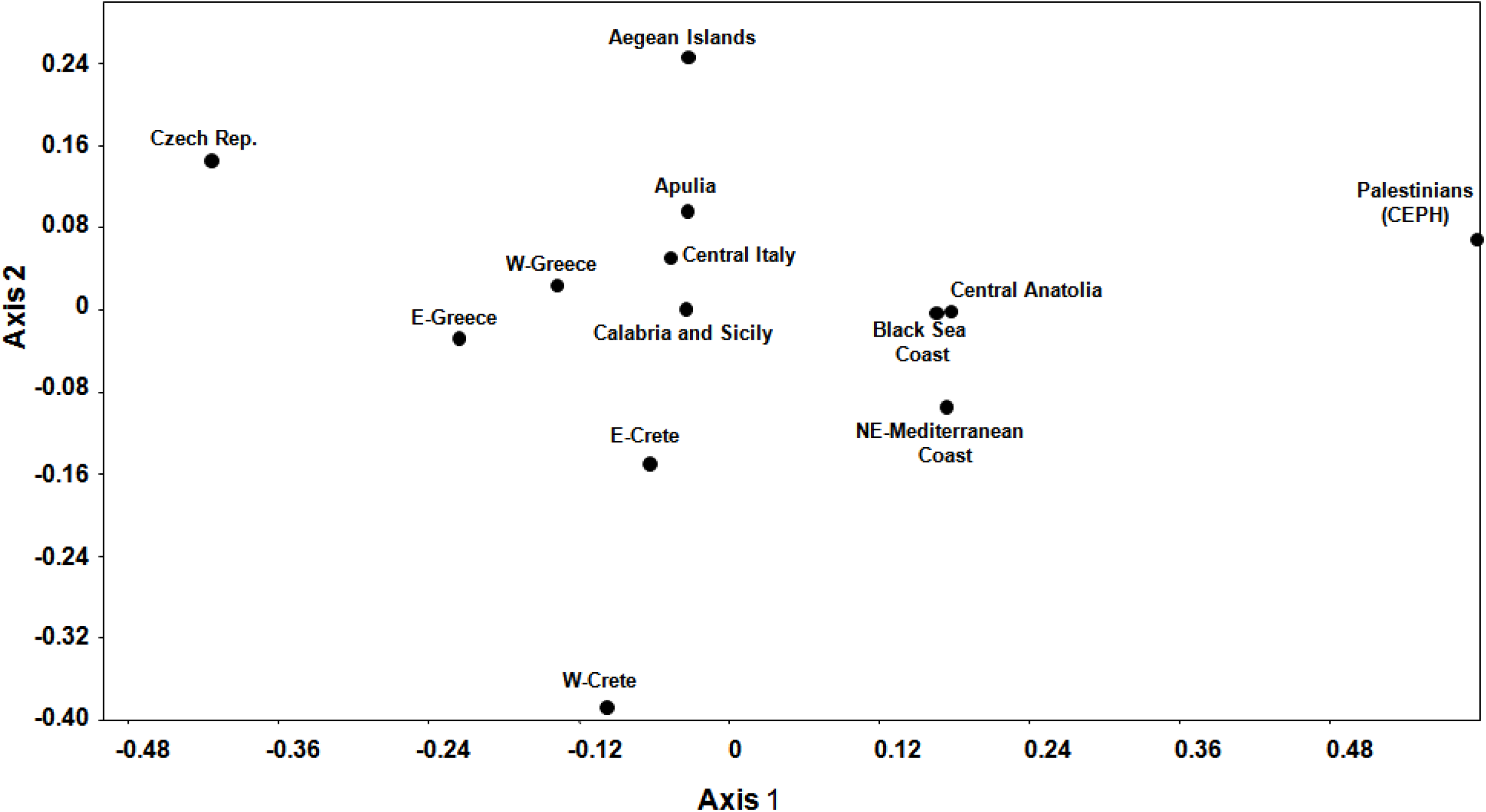
Scatterplot of the scores on the first two axes obtained by nmMDS based on the matrix of pairwise Fst values after grouping the 41 location samples into 13 geographical pools

**Supplemental Fig. 6.**
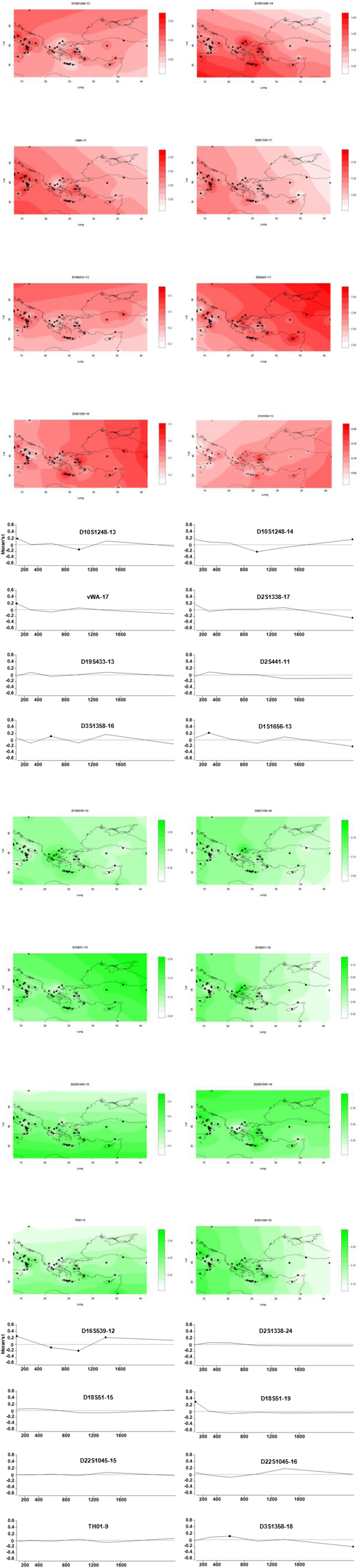
Frequency contour maps of 8 alleles producing the highest squared loadings on sPC 1 (page 1, red) and sPC 2 (page 3, green). The same alleles are listed in Table 1 in the 2.5% columns. Note the different colour scale used in each map. Values outside the polygon connecting the most external points are extrapolated. For each of the two map sets, the correlograms are shown in the same order (pages 2, 4). Black dots indicate significant class-specific values. Ticks on the x axis are spaced to indicate the upper bounds of distance classes.

**Supplemental Fig. 7.**
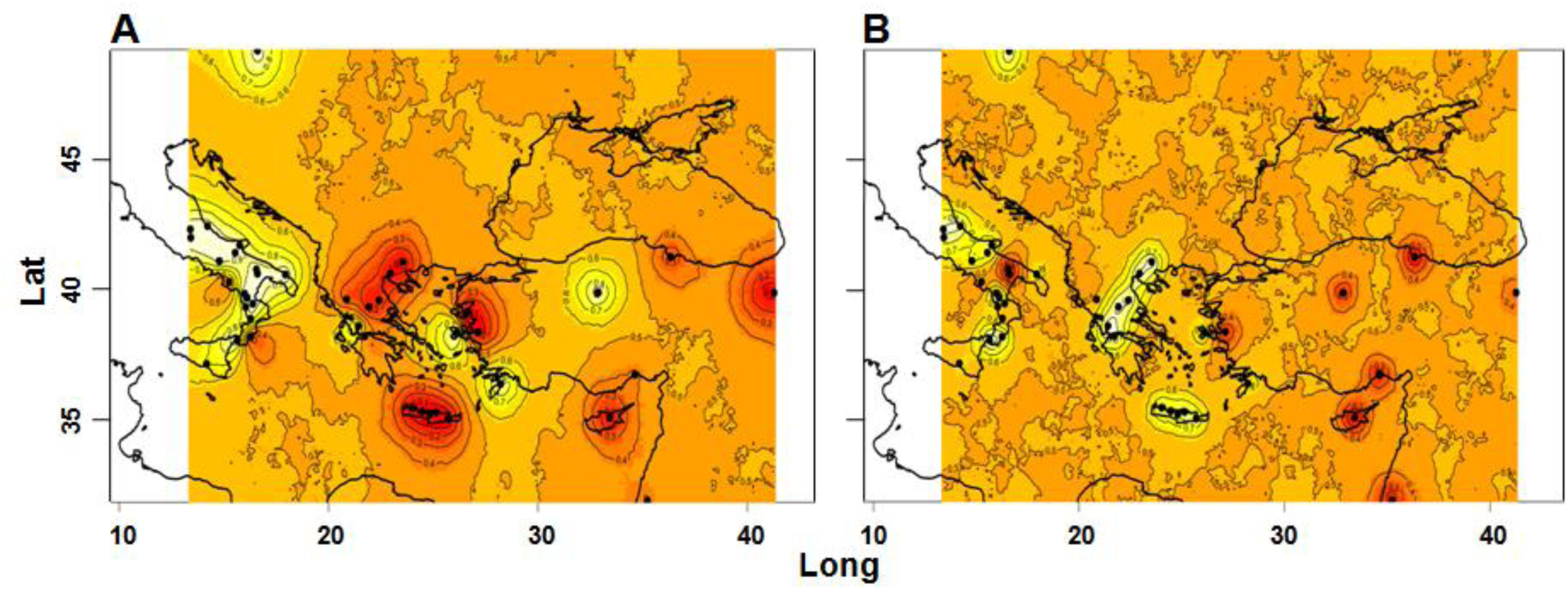
Color maps (corresponding to Figure 1C,D) of assignment probabilities of the 41 locations to either of 2 population clusters obtained with Geneland on the reduced allele sets derived from sPC1 (A) and sPC2 (B).

**Supplemental Fig. 8.**
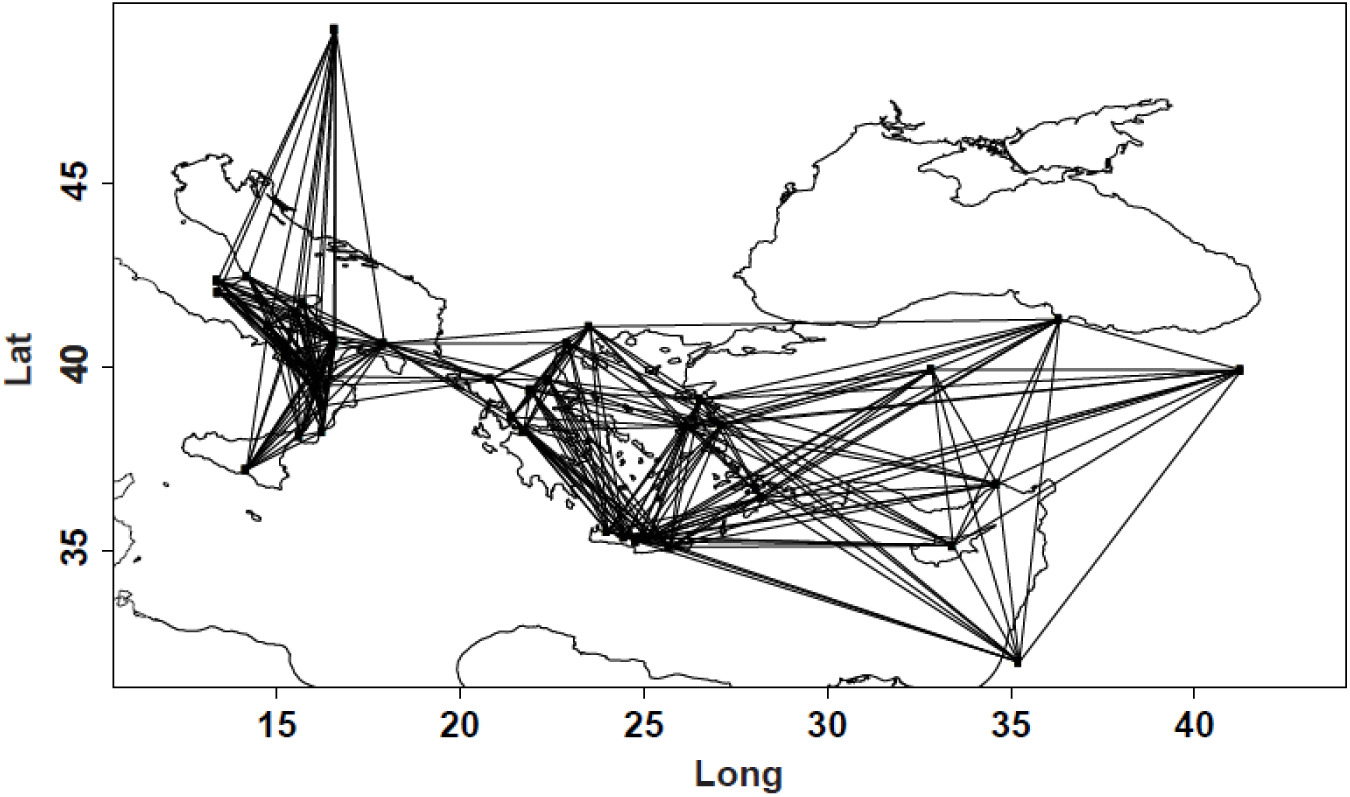
Map of Southern Europe/Northeastern Mediterranean Sea, with the nearest neighbour (n=12) connection network used in adegenet.

**Supplemental Fig. 9.**
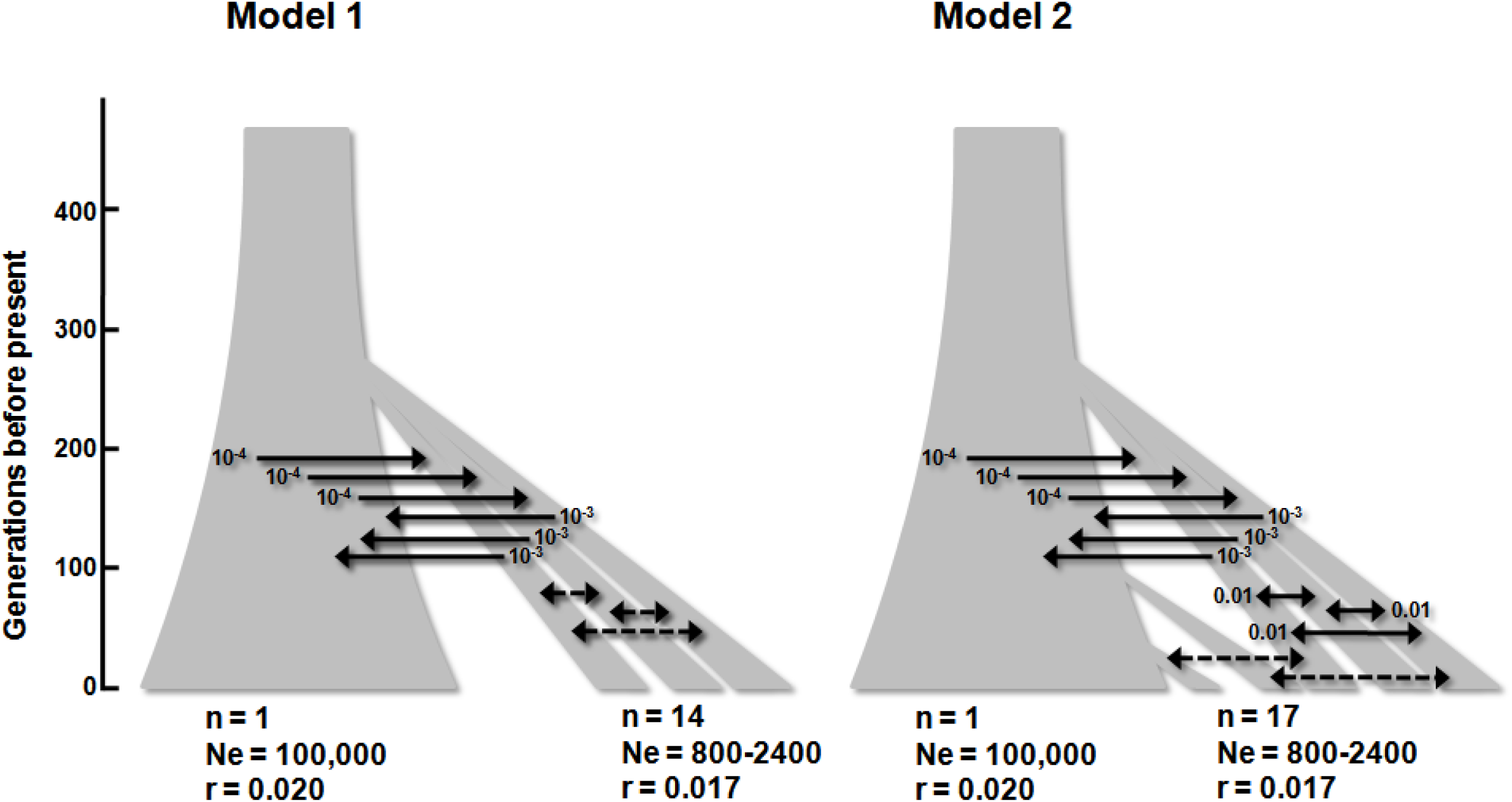
Scheme of the demographic models used for coalescent simulations. n = number of demes (only a subset shown for the sake of clarity); Ne = effective size (in gene copies); r = growth rate per generation. Black arrows represent instances of gene flow with the indicated fixed rate across simulations. Dashed arrows indicate instances of gene flow whose rate was varied across simulations.

**Supplemental Fig. 10.**
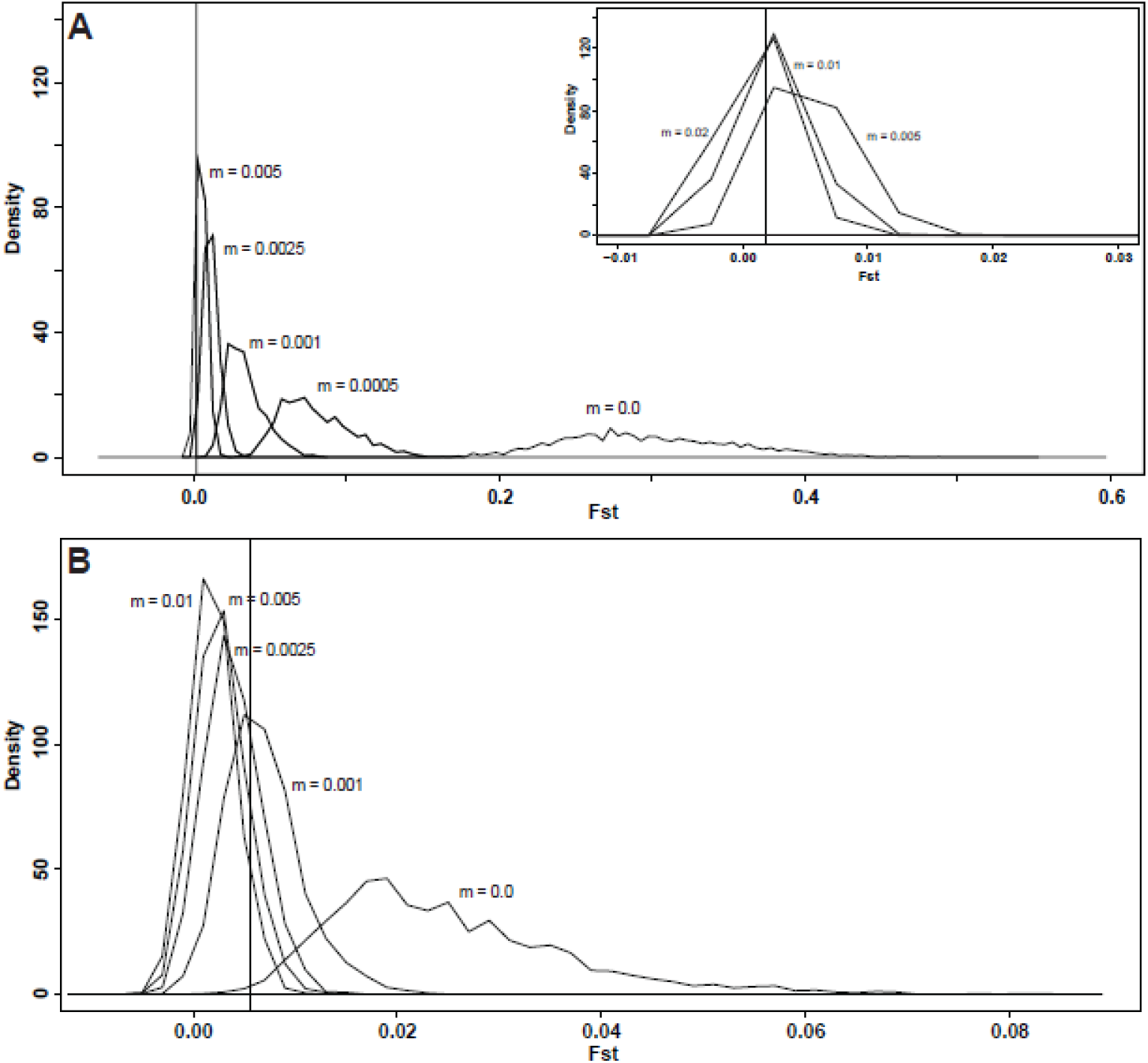
Distributions of simulated Fst values under the demographic scenarios of Supplemental Fig. 3. A) Model 1: migration rates (m) among the 14 demes are shown; the curves for m=0.02, 0.01 and 0.005 are shown in the inset for clarity. B) Model 2: the numbers indicate migration rates (m) between 3 recent demes and the 14 demes of Model 2. Note the different scale of the x axis as compared to panel A. In both panels the vertical lines indicate the Fst value obtained from real data.

**Supplemental Table 1.** List of sampling locations, geographic coordinates and sample sizes. For Italy and Greece the administrative region or the Island name is also reported.

|  | Sampling location | Country/Island | Long E | Lat N | Sample size (n. of subjects) |
| --- | --- | --- | --- | --- | --- |
| 1 | Brno | Czech Rep. (Moravia) | 16.607 | 49.195 | 49 |
| 2 | L'Aquila | Italy (Abruzzo) | 13.400 | 42.350 | 32 |
| 3 | Avezzano | Italy (Abruzzo) | 13.426 | 42.028 | 28 |
| 4 | Pescara | Italy (Abruzzo) | 14.216 | 42.462 | 18 |
| 5 | Benevento | Italy (Campania) | 14.783 | 41.130 | 45 |
| 6 | Foggia | Italy (Apulia) | 15.545 | 41.462 | 26 |
| 7 | Gargano promontory | Italy (Apulia) | 15.750 | 41.733 | 30 |
| 8 | Cilento promontory | Italy (Campania) | 15.250 | 40.333 | 44 |
| 9 | Matera | Italy (Basilicata) | 16.604 | 40.666 | 34 |
| 10 | Altamura | Italy (Apulia) | 16.553 | 40.825 | 20 |
| 11 | Brindisi | Italy (Apulia) | 17.942 | 40.633 | 93 |
| 12 | Lungro | Italy (Calabria) | 16.123 | 39.742 | 24 |
| 13 | Acri | Italy (Calabria) | 16.383 | 39.489 | 30 |
| 14 | Mormanno | Italy (Calabria) | 15.986 | 39.890 | 36 |
| 15 | Paola | Italy (Calabria) | 16.041 | 39.360 | 38 |
| 16 | Lamezia | Italy (Calabria) | 16.276 | 38.934 | 17 |
| 17 | Locri | Italy (Calabria) | 16.264 | 38.234 | 38 |
| 18 | Reggio Calabria | Italy (Calabria) | 15.647 | 38.111 | 41 |
| 19 | Butera | Italy (Sicily) | 14.182 | 37.192 | 24 |
| 20 | Ioannina | Greece (Epirus) | 20.854 | 39.665 | 25 |
| 21 | Agrinion | Greece (Aetolia-Acarnania) | 21.410 | 38.624 | 22 |
| 22 | Patrai | Greece (Akhaia) | 21.735 | 38.247 | 62 |
| 23 | Kardhitsa | Greece (Thessaly) | 21.921 | 39.364 | 17 |
| 24 | Larisa | Greece (Thessaly) | 22.419 | 39.639 | 17 |
| 25 | Thessaloniki | Greece (Central Macedonia) | 22.944 | 40.640 | 15 |
| 26 | Serrai | Greece (Central Macedonia) | 23.541 | 41.091 | 27 |
| 27 | Mitilini | Greece (Lesvos) | 26.557 | 39.107 | 18 |
| 28 | Khios | Greece (Khios) | 26.136 | 38.371 | 30 |
| 29 | Rhodos | Greece (Rhodos) | 28.217 | 36.435 | 50 |
| 30 | Crete (unspecified) | Greece (Crete) | 24.809 | 35.240 | 65 |
| 31 | Khania | Greece (Crete) | 24.018 | 35.514 | 45 |
| 32 | Rethmnon | Greece (Crete) | 24.482 | 35.364 | 67 |
| 33 | Iraklion | Greece (Crete) | 25.144 | 35.339 | 48 |
| 34 | Lasithi plateau | Greece (Crete) | 25.710 | 35.081 | 38 |
| 35 | Western Mediterranean coast | Turkey | 27.129 | 38.419 | 54 |
| 36 | Central Anatolia | Turkey | 32.854 | 39.921 | 94 |
| 37 | Black Sea coast | Turkey | 36.331 | 41.293 | 42 |
| 38 | Eastern Anatolia | Turkey | 41.277 | 39.909 | 33 |
| 39 | Eastern Mediterranean coast | Turkey | 34.633 | 36.800 | 27 |
| 40 | Cyprus | Turkey (Cyprus) | 33.430 | 35.126 | 46 |
| 41 | Palestine (CEPH) | Palestine | 35.233 | 31.952 | 50 |

**Supplemental Table 2.** List of alleles not matching the AmpFLSTR^®^ NGM SElect^™^ allelic ladder (overladder). Loci are in the order of increasing MW for the Blue, Green, Black and Red series.

| Locus | Allele | n. of allele<br>copies | Reported in<br>http://www.cstl.nist.gov/<br>YES/NO |
| --- | --- | --- | --- |
| D10s1248 | None |  |  |
| vWA | None |  |  |
| D16S539 | None |  |  |
| D2S1338 | 13 | 2 | Y |
| D8S1179 | 7.3 | 1 | N |
| D8S1179 | 10.3 | 1 | Y |
| D2S11 | 28.1 | 1 | Y |
| D2S11 | 29.1 | 3 | Y |
| D2S11 | 30.1 | 3 | Y |
| D2S11 | 31.3 | 1 | Y |
| D18S51 | 22.1 | 1 | Y |
| D22S1045 | None |  |  |
| D19S433 | 6.2 | 2 | Y |
| D19S433 | 7 | 1 | Y |
| D19S433 | 19.2 | 1 | Y |
| TH01 | None |  |  |
| FGA | 19.1 | 1 | Y |
| D2S441 | None |  |  |
| D3S1358 | None |  |  |
| D1S1656 | 8 | 1 | Y |
| D12S391 | 17.1 | 1 | Y |
| D12S391 | 18.2 | 1 | Y |
| D12S391 | 19.1 | 1 | Y |
| D12S391 | 21.3 | 2 | Y |
| SE33 | 7.3 | 1 | Y |
| SE33 | 13.3 | 2 | N |
| SE33 | 14.1 | 1 | N |
| SE33 | 14.3 | 5 | N |
| SE33 | 15.3 | 4 | Y |
| SE33 | 17.3 | 2 | Y |
| SE33 | 18.3 | 3 | Y |
| SE33 | 25.3 | 1 | N |
| SE33 | 27.3 | 1 | N |
| SE33 | 30.1 | 1 | N |
| SE33 | 39 | 1 | N |

Supplemental Table 3. Spreadsheet with relative allele frequencies at 16 STR loci in the 41 location samples.

This large table is available upon request to A. N.

